# Autonomous induction of hepatic polarity to construct single cell liver

**DOI:** 10.1101/636654

**Authors:** Yue Zhang, Richard de Mets, Cornelia Monzel, Pearlyn Toh, Noemi Van Hul, Soon Seng Ng, S. Tamir Rashid, Virgile Viasnoff

## Abstract

Symmetry breaking of protein distribution and cytoskeleton organization is an essential aspect for development of apico-basal polarity. In embryonic cells this process is largely cell autonomous, while differentiated epithelial cells collectively polarize during epithelium formation. We report here that the *de novo* polarization of mature hepatocytes is a cell autonomous process. Single hepatocytes developed *bona fide* secretory hemi-apical lumens upon adhesion to finely tuned substrates bio-functionalized with cadherin and extra cellular matrix. The creation of this single cell liver allows unprecedented control and imaging resolution of the lumenogenesis process. We demonstrate that the density and localization of cadherins along the initial cell-cell contact acted as a key factor triggering the reorganization from lateral to apical actin cortex. Consequently, we established why hepatocytes could form asymmetric lumens in heterotypic doublets involving another ectopic epithelial cell originating from kidney, breast, or colon.

## Introduction

The development of apico-basal polarity in epithelial cells requires signaling from the extracellular matrix as well as from cell-cell contact. The matrix/cell junction direction provides an axis of external cues along which epithelial cells break their symmetry (*1–3*), organize their acto-myosin cortex (*4*), and direct the vectorial transport of proteins (*5, 6*). Mechanical constrains and tension also play a role (*7–9*). The segregation of membrane proteins between apical, basal, and lateral poles are also ensured by the fencing activity of tight junctions as well as by the antagonist pathways of the polarity complexes (Par1/Par2, Par3/Par6/aPKC, Crumbs and Scribble complexes). The situation differs in the early embryo when single cells can polarize in the absence of external cues. In single cell *C*.*elegans* embryo, Par 1 and Par 3 complexes segregate based on antagonist kinase activity (*10, 11*) and acto-myosin cortical flow (*12*). In mouse embryo, blastocysts develop apical poles (*13, 14*) in the absence of extracellular matrix interactions in a fully cell autonomous manner (*15, 16*). Over activation of LKB1 in single epithelial cells was also reported to stimulate the creation of a border brush with a partial localization of the polarity complexes (*17*). The question then arises whether *de novo* establishment of epithelial polarity is a cell autonomous response triggered by external cues or a collective response associated to the concomitant development of polarity in neighboring cells. Usual epithelial models (tissue, cell monolayer, or cysts) inherently fail to address this question. Considering a cell polarizing in these multi-cellular contexts, the concomitant reorganization of cell-cell contacts of the neighboring cells can act as a polarization cue and as a response to its polarization establishment.

This study presents a novel model where single primary hepatocytes develop independent *bona fide* secretory apical poles when grown in synthetic microenvironments. In this context, we demonstrate for the first time that *de novo* apical lumen development is a genetically controlled cell autonomous process. It depends mainly on actin cortex rearrangements triggered by the biophysical properties of the cadherin-mediated adherens junction along the initial lateral cell-cell contact. Subsequently, we demonstrate that, hepatocytes can form functional lumen with a whole variety of epithelial cells. In particular, we show that mature hepatocytes can polarize with immature hepatocytes during the differentiation process.

## Results

We investigated the *de novo* lumenogenesis of bile canaliculi to test if the apical polarity development is a cell autonomous process triggered by simple cues sensed by the cell along the non-polarized initial cell-cell contact. We used hepatocytes at different maturation stages during differentiation. Mature hepatocytes develop intercellular secretory apical lumen called bile canaliculi, connecting the hepatocytes to the biliary system. They consist of small, elongated tubules (2 μm in diameter) sealed by tight junctions and extending between two adjacent cells (**Figure 1a**). Bile salt transporters (eg:BSEP) and ion pumps (eg:MRP2) accumulate at the apical membrane to secrete bile into the lumen. We differentiated hiPSCs (human induced pluripotent stem cells) over 25 days (*18, 19*) into hepatocytes lineage during which the hepatoblast-to-hepatocyte differentiation took 6 days in vitro (*20*). During this hepatic lineage induction, we observed a mixed population of cells at different maturation stages (*21*). To assess hepatocyte maturity, we used the expression levels and localization of a small organic anion transporter MRP2 (Multidrug resistance-associated protein 2). A mixed population of mature and immature hepatocytes was plated in micro-fabricated cavities bio-functionalized with fibronectin (see protocol). We previously established that such cavities forced cell-cell contacts to enable lumen formation (*9*). We found that homotypic junction between two mature, MRP2 positive hepatocytes expectedly developed a canaliculus (**Figure 1a**). The lumen displayed a symmetric accumulation of MRP2 exclusively at their apical membrane. The apical pole from both cells displayed characteristic actin enrichment. The lumens were inflated due to the proper localization of tight junctions at their edges (**Figure 1a**). We also observed a sizeable fraction (40%±5) of heterotypic junctions between mature and immature hepatocytes (MRP2^+^ / MRP2^−^ cells). They displayed strikingly asymmetric apical lumens (**Figure 1a**). The luminal membrane of the mature hepatocyte (MRP2^+^) was identical to the canaliculus formed in homotypic junctions between mature hepatocytes. By contrast, the membrane of the immature hepatocyte (MRP2^−^) lacked actin enrichment and was hardly curved. However, tight junctions, as shown by the presence of ZO1, properly sealed the edges of the asymmetric lumens (**Figure 1a** and **Supplementary Figure 1a**). The asymmetric apical arrangement of canaliculi suggested that the nature of the cell adjacent to a mature hepatocyte might not be critical for the hepatocyte to establish apical basal polarity. To further test this hypothesis, we induced heterotypic contacts between primary rat hepatocytes (*22, 23*) and epithelial cell lines derived from various species and organs: EPH4, a murine breast cell line, Madin Darby Canine Kidney cells (MDCK), a dog kidney cell line and Caco2, a human colorectal adenocarcinoma cell line. Monolayer co-cultures of these cell lines lead to spontaneous segregation of the population. However, constraining the cells in microcavities (25×25×25μm) favored the establishment of stable heterotypic contacts. A large number of lumens formed along these heterotypic contacts. They presented the proper localization of the respective apical markers for each cell type (MRP2 for the hepatocytes, GBP35 for MDCK, **Figure 1b** and **Supplementary Figure 1b**). All luminal membranes exhibited microvilli. ZO1 also localized at the lumen edge indicating a normal polarized state for both cells. The large inflation of the lumen revealed an efficient paracellular barrier and the development of transluminal osmotic gradients.

**Figure 1:**
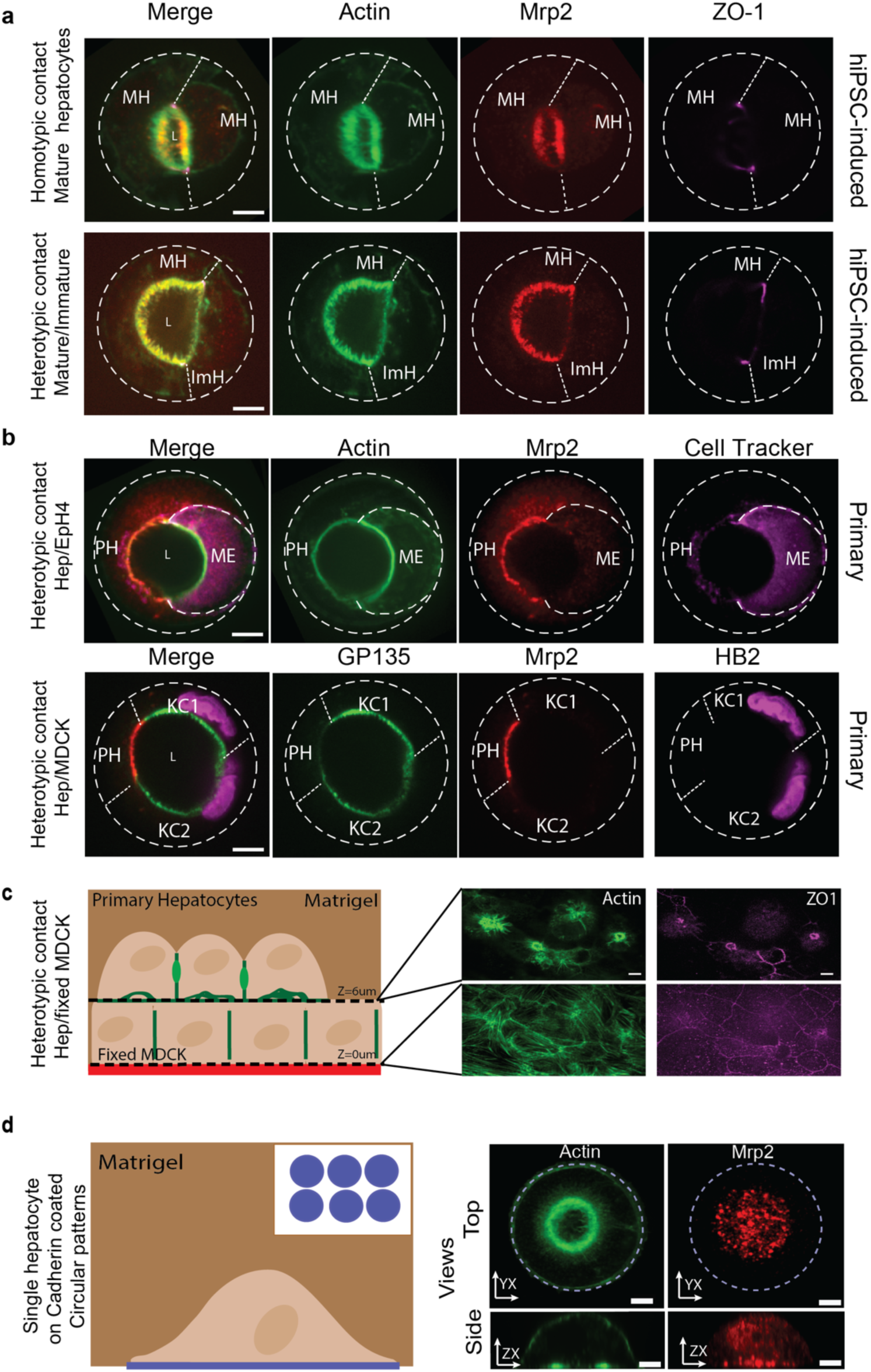
Hepatocytes are able to form lumens with different types of substrates. **a**, Overlay of confocal images of the lumen (L) created inside a microwell between mature (MH) and immature hepatocytes (ImH) stained for Actin (green), Mrp2 (red) and ZO-1 (magenta) respectively. The well wall and cell junction has been delimited by dashed lines. Scale bar = 10μm. **b**, Representative confocal images of the lumen (L) created inside a microwell between primary hepatocyte (PH) and different cell lines such as mammary epithelial cells (Eph4, ME) or kidney cells (MDCK, KC) stained with Mrp2 (red), actin/golgi (green) and cell tracker/histones (magenta). Scale bar = 10μm. **c**, Left, side view illustration of the co-culture of fixed MDCK with primary hepatocytes. Right, confocal images of actin (green) and ZO-1 circular structures (magenta) at the interface between primary hepatocytes and fixed MDCK (Z=6μm) after 24h of seeding, and between fixed MDCK with the coverslip (Z=0μm). Scale bar = 10μm. **d**, Left, Side view illustration of single cell primary hepatocytes. Right, Immunostaining of actin (green) and Mrp2 (red) of primary hepatocytes on an E-cadherin island covered with matrigel reveal an actin ring located at the interface with the coverslip and a preferential localisation of Mrp2 above the actin ring. Scale bar = 10μm.

This result demonstrated that the development of interspecific lumens occurs in the absence of inhibitory mechanisms. It supports the hypothesis that apico-basal polarity develops stably upon response to signaling cues ubiquitous to epithelial cell types. In the heterotypic doublets, the organization of the tight junctions was nonetheless symmetrical. We thus could not decipher if the polarity is triggered autonomously based on cues present at the initial cell-cell contact, or if it requires the concomitant organization of both cells in a coordinated manner.

We then tested if single hepatocytes could autonomously polarize in absence of polarizing neighboring cells. To this end, we first cultured and fixed (paraformaldehyde 4%) MDCK monolayers at 65% confluence and used them as an “inert” substrate to culture fresh primary rat hepatocytes (Material and Methods). A substantial fraction of single hepatocytes (~50%) developed unusual accumulation of actin filaments or actin rings at the interface between the MDCK monolayer and the hepatocytes **(Figure 1c)**. ZO1 accumulated around these actin rings. We reasoned that these structures could constitute the basis of a polarized domain that self-organized upon the engagement of hepatocytes in an E-cadherin mediated contact with the fixed MDCK.

To test this hypothesis we reconstituted cell-cell and cell-matrix interactions *in vitro* using biofunctionalized substrates enabling precise control of the interaction and high-resolution imaging. We screened various environmental conditions (**Supplementary Figure 2a**). On 700µm^2^ circular islands coated with fibronectin, single hepatocytes spread and organized their actin cytoskeleton around the pattern showing no sign of polarity, and this irrespective of whether the culture medium was supplemented with 6% Matrigel. On identical islands coated with E or N cadherins, single cells developed central actin rings at the interface with the substrate only for Matrigel supplemented medium (**Supplementary Figure 2b**). Note that Hepatocytes express equally E and N-Cadherins (*24*). MRP2 was recruited within the membrane area delineated by the actin ring. Similar results were obtained for primary mouse hepatocytes (**Supplementary Figure 2c**).

We further tested if the inner ring membrane was a functional secretory apical pole as suggested by the localization of the anion transporter MRP2. Confocal imaging did not reveal any convincing detachment of the membrane from the substrate suggesting either a hemi-luminal cavity below optical axial resolution (around 700nm) or a collapse during fixation. We thus used Reflection Interference Contrast Microscopy (RICM) to measure the nanoscale distance between the substrate and the plasma membrane in label-free living cells (*25*). This revealed (**Figure 2 a**) that the membrane enclosed by the actin rings had a concave shape on average. The membrane pulsated up and down at mean period of 6min/pulse. (N=9). The hemi-lumen reached maximal heights of 170±5 nm (N=25). Upon partial inhibition of bile salt synthesis by 10µm Ketoconazole, the lumen period increased by to fold to 12 min/pulse. It also enhanced lateral fluctuations of the intraluminal membrane as compared to the concentric axial pulsation observed in control case (**Figure 2a** and **Supplementary movies 1-2**). Additionally, actin filaments no longer formed a characteristic ring but remained structured as a patch in the center of the contact. We also added UDCA (Ursodeoxycholic acid, 40µM) to the cell culture media to stimulate bile secretion, which resulted in three-fold increase (633±54nm N=16) in lumen maximal height. The central lumen was largely inflated as demonstrated by the multiple circular interference fringes along the lumen. Our data strongly suggested that the hemi-lumens formed between hepatocyte and cadherin-coated micropattern were functional and recapitulated the pulsatile behavior of canaliculi observed *in vivo*. The small level of inflation of the lumen (around 200nm as compared to 2 to 5 μm *in vivo*) could be explained by the expected large paracellular leak resulting from a contact with the substrate that is established exclusively by cadherins and likely lacking tight junctions. We then tested the extent to which the spatial self-organization of junctional and polarity markers resulting solely from E-cadherin adhesion mimicked the organization of true canaliculi. **Figure 2b** shows the positions of cellular cadherin, ZO1/ZO2, Par-3, Claudin-3, Myosin IIA, MRP2 relative to the actin ring. The stereotypical shape of the ring enabled the computation of the average localization map (**Figure 2c and Supplementary Figure 3**) for each protein (**Material and Methods**). From this map we found that myosin II microfilaments accumulated strongly along the ring. Structured illumination microscopy (**Supplementary Figure 7**) revealed their radial orientation across the orthoradial actin fibers constituting the ring, strongly suggesting that the ring is highly contractile. ZO1/ZO2 and Par3 accumulated precisely at the edge of the actin ring (**Figure 2b,c**). Cellular cadherin (**Material and Methods**) was absent from the central region of the ring, accumulated 1.5 ± 0. 5 μm away from the outer edge of the ZO1-ZO2-Par3 ring and was diffusively present along the rest of the contact with the substrate. We concluded that proper lateral, junctional, and basal domains developed at the contact with the substrate as pictured on **Figure 2d**. Note however that the different concentric rings extend laterally over 2 ± 0. 5 μm. In canaliculi formed between 2 cells, the peri-canalicular acto-myosin ring was smaller than 2 microns, and ZOs and Par3 localized around a 200-300 nm away from the actin belts (*26*). Additionally, the main trans-membrane tight junction proteins claudin3 and 1 and occludin did not show specific accumulation on any membrane region, indicating a lack of structuration of the transmembrane components of the tight junction components (**Supplementary Figure 3a**). The minimalistic cues provided by our system not only polarized the cortical and membrane components at the contact but also structured the position of the Golgi. 3D reconstitution following Grasp65 staining (**Supplementary Figure 3b**) revealed that the Golgi was located precisely (95% overlap N=40, **Material and Methods**) over the actin ring region. It extended vertically to reach the nucleus independently of the position of the nucleus in the cells. Although it was previously reported that cadherin and integrins are necessary to elicit the development of apical poles (*2, 3*), in our model the polarity developed without cell division (as in classical type of approaches (*27*)) and by a single cell in contact with an inert substrate. Taken together, we established that combining static cadherin and extracellular matrix adhesions is sufficient to induce a fully polarized protein distribution which phenocopied the apico-basal polarization of hepatocytes (except for occludin and claudins) (**Figure 2d**). Our data demonstrated that polarity program in mature hepatocytes is fully autonomous and does not require the response of the neighboring cells past the initial induction by E-cadherin contact.

**Figure 2:**
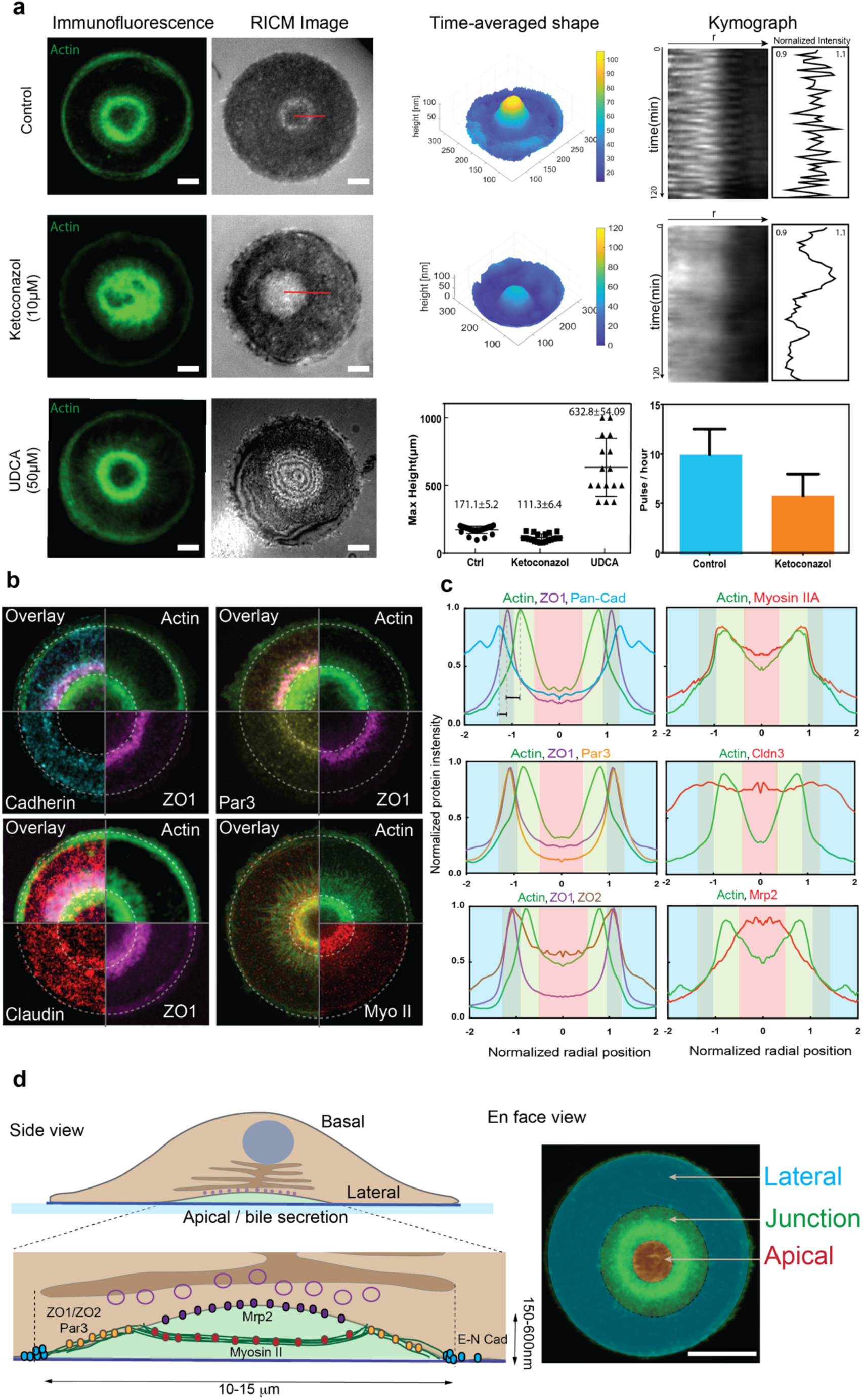
Characterisation of protein organization and behaviour of single-cell hemi-lumen. **a**, **Left**, immunostaining and RICM images of single-cell liver following different drug treatments. **Right**, quantification of height and pulsation behaviour under the three different treatments. Reduction of bile secretion by ketoconazole induces a homogeneous bright circle that is smaller, and with a slow pulsation compared to control. Boosting secretion by UDCA induces an inflation of the lumen resulting in multiple interference rings on the RICM images. N_ctrl_=25, N_ket_=20, N_UDCA_=16, Scale bar = 10μm. **b** and **c**, Montage and quantification of confocal images of the hemi-lumen stained for structural and polarity markers illustrates the spatial localisation of the different rings of proteins around the lumen. For quantification, the centre of the lumen is considered as 0 while the edge of the actin ring is considered as 1. The red, dark green, light green and blue background correspond to the region described in d. **d**, Side and en face view illustration of the position of the different proteins studied and discretisation of the cell-coverslip interface in four regions. The apical pole (red region) is delimited by the actin ring (light green region). As well as containing actin, this inner ring is rich in myosin IIA. Moving outwards, rings of ZO1/ZO2 and Par3 are found (dark green region, followed by cell-cell contacts labelled by E-cadherin (blue region). E-cadherin is not observed from the centre of the lumen out to the ZO1/ZO2 ring. Mrp2 is mostly located above the apical pole. In this system, claudins did not exhibit any specific localisation. Scale bar = 20μm, N_ActZO1Cad_=9, N_ActMyosin_=14, N_ActZO1Par3_=17, N_ActCldn3_=10, N_ActZO1ZO2_=5, N_ActMrp2_=11.

We then established the time sequence of events leading to the development of apico-basal polarity after it was triggered by mere E-cadherin contact. We classified the development of the hemi-lumen into five phases based on the different prevailing acto-myosin structures imaged by structured illumination microscopy, SIM (**Figure 3**).

**Figure 3:**
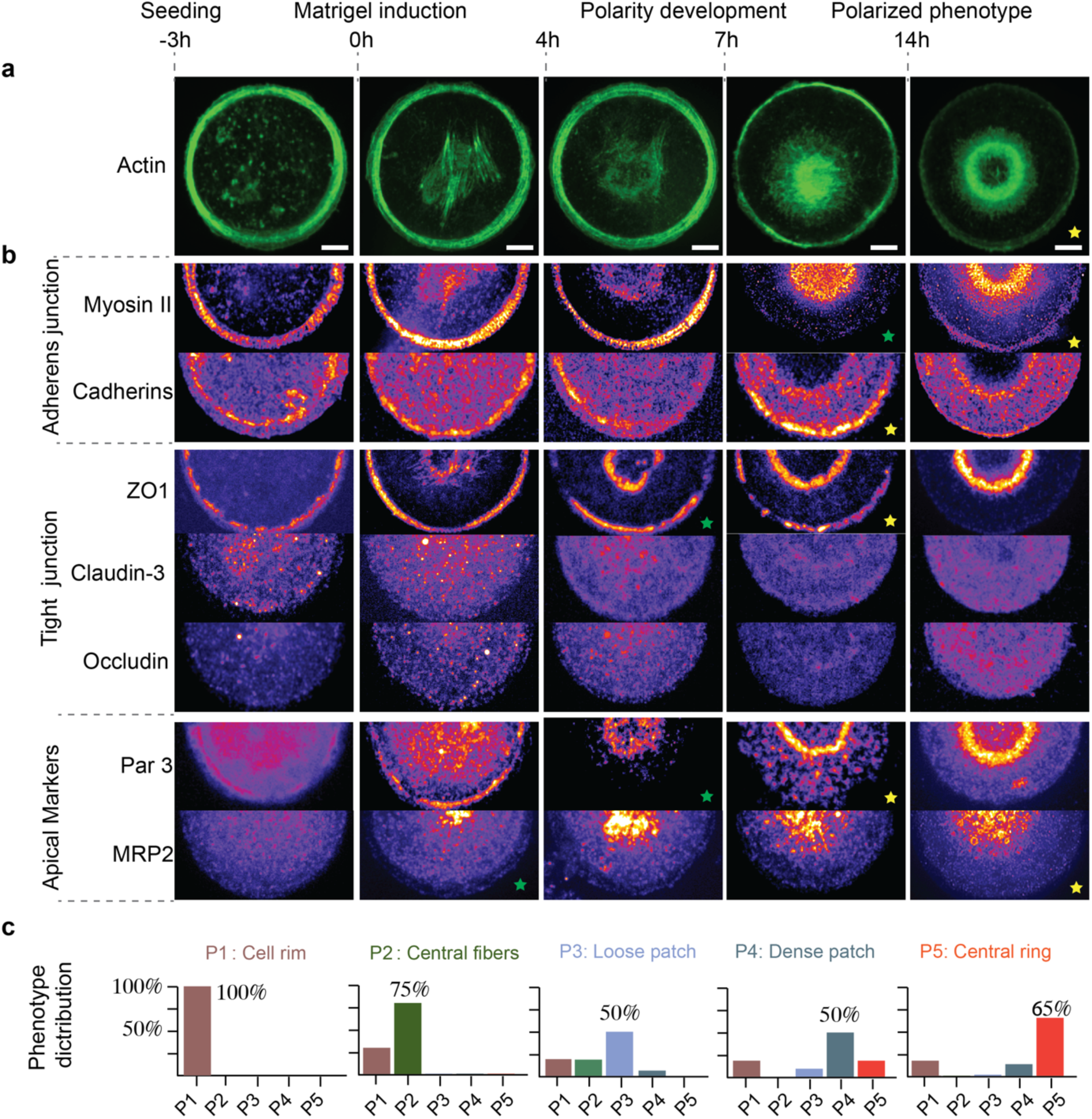
Actin and hepatic polarity related proteins undergo extensive reorganization throughout polarity establishment. **a,** Representative Structured Illumination Microscopy (SIM) images showing the formation of an actin ring around the apical pole of single hepatocytes situated on E-cadherin circular patterns fixed at 3 hours before, and then 0, 4, 7, 10 and 14 hours after matrigel addition. Scale bar, 5µm. **b,** Representative SIM images of adherens junction associated proteins (Cadherins and Myosin II), cytosolic and transmembrane components of tight junction (ZO-1, claudin-3 and, occludin), apical markers (Par3 and MRP2) at five stages of polarity development. **c,** Fraction of cells displaying typical phenotypes of each development phase fixed at different time points. Based on the actin organization, five phases are defined as indicated. The number of cells analyzed was pooled from n = 3 independent experiments (N= 42 for P1, N= 65 for P2, N= 56 for P3, N= 66 for P4, N= 72 for P5).

### Phase 1: 3h pre matrigel induction

Cells first adhered to the E-cad patterns. During this phase an acto-myosin ring accumulated at the cell edge. Cadherins and ZO1 co-localized with the ring. Other markers showed no specific localization.

### Phase 2: 0h to 4h post induction

Disorganized actin fibrils developed at the center of the cell contact with the substrate. Myosin IIA and ZO-1 were recruited along these fibers. Par3 accumulated diffusively in the central region.

### Phase 3: 4h to 7h post induction

A diffuse acto-myosin patch developed in the center of the cells (50% cases, N=56). ZO1/2 accumulated at the outer edge of the patch into a discontinuous ring. Par3 and MRP2 localized diffusively over the patch. Cadherin attachment to the substrate remained homogenous across the whole contact area.

### Phase 4: 7h to 14h post induction

The acto-myosin patch densified into a disorganized central region surrounded by radial fibers. Myosin was largely recruited on the disorganized patch (**Supplementary Figure 7**). Cadherin detached from the underlying substrate beneath the central patch region. Polarity markers (ZO1-2, Par3) were densely recruited around the interface between the patch and the radial fibers. MRP2 accumulated within the central patch region.

### Phase 5: 14h post induction

Hepatocytes developed the ring phenotype described previously (**Figure 2**) with 85% occurrence rate (N=72). Note that ZO1/2 were fully excluded from the edge of the cell. The transition from Phase 4 to Phase 5 was suppressed by inhibition of myosin II or bile secretion (**Supplementary Figure 4**), strongly suggesting that the transition from actin patch to actin ring was mechanically driven by acto-myosin contractility and osmotic gradients.

We then tested if this time sequence of events is compatible with the development of real lumens between two cells. We performed 3D SIM imaging of different development stages on the canaliculi (**Supplementary Figure 5**). They displayed gradual accumulation of a denser actin cortex (300 nm thick) at the center of the contact and the gradual relocalization of ZO1 and Par 3 from the contact edge to the lumen edge. Lumen inflation occurred in multiple locations along the actin dense patch. We attributed this effect to the maturation of tight junctions concomitant to the disengagement of adherens junctions. This leads to the local development of micro lumens that eventually merged into one lumen. We concluded that the process of canaliculi development *in vivo* and in our reductionist single cell liver follow a very similar sequence of events. The time sequence of events we observed also share similarities with what has been described as pre-apical actin patches (PAP) and Apical Membrane Initiation Sites (AMIS) (*4, 28*) in MDCK doublets. However, in our case the actin structure formed *in vivo* is far more localized at the membrane than what is reported for MDCK PAP (*4*).

We then determined how the density and spatial arrangement of cadherin regulates lumen formation. We first modulated the total amount of cadherin adhesion by changing the shape and size of the pattern while keeping cadherin density constant. Independent of the pattern’s shape and size (**Supplementary Figure 6**), lumens formed with identical rate of occurrence, and remained circular with an area of 200 μm^2^ ± 60. Unconstrained hepatocytes left to spread on un-patterned homogeneously coated cadherin substrates also polarized. These lumens were more irregular in shape, and could reach an area of 600 μm^2^ for very large contacts (2400 μm^2^). We concluded that in our reductionist approach the lumen size and shape were largely decoupled from the size and shape of the cadherin contact area.

Next, we varied cadherin density on the 30 μm circular pattern (**Material and methods**). **Figure 4a** shows that at low cadherin densities the hepatocytes did not attach. As the cadherin density increase the number of hepatocytes adhering to the substrate continuously increased, however the occurrence of lumen formation peaked significantly at a coating density of 10 μg/ml cadherin. Lower and higher cadherin densities proved less efficient in prompting lumen formation suggesting that there is an optimal density of cadherin for triggering apical surface development. We used western blotting to assess the quantity of cadherins. In ascending expression levels, the cell lines ranked as MDCK < EPh4 < Caco2 (**Supplementary Figure 1b**). In the heterodoublets the probability of asymmetric lumens formation was inversely correlated with the amount of cadherin. MDCK cells had the lowest expression levels of cadherin and proved the most efficient in creating heterolumens. We then overexpressed E-cadherin in Eph4 cells (Eph4^+^) to reach the expression level found in Caco2 cells. It resulted in a lower occurrence of lumen formation matching that matching Caco2 heterodoublets (**Figure 4a**). We thus concluded that apical pole formation requires a fine balance of cadherin adhesion. On one hand it should allow cell-cell contact and on the other hand should not over-stabilize it. This parameter appears to be key for the ability of hepatocytes to develop mature lumens with their adjacent neighbors. We then tested if the spatial distribution of cadherin also affected lumen formation. We seeded our single hepatocytes on circular cadherin patterns (30 μm ⃠) containing an antifouling coating in their central region (15 μm ⃠). On the center of these doughnut patterns, we matched the size of the non-adhesive region to the average size of lumens formed on disc patterns (**Figure 4b**). All these hepatocytes failed to form a secretory hemi-lumen, and the membranes remained suspended over the non-adhesive part of the lumen as demonstrated by the random and stochastic fluctuations probed by RICM live imaging (**Supplementary Video 4**). On average, this resulted in a flat top-hat profile of the membrane height as compared to the dome shape profile observed when lumens formed (**Figure 4b**). The actin structure remained cortical with no radial fibers (**Figure 4c**), and never developed into a ring. ZO-1 and Par-3 relocated to the central zone but did not organize along the contour of actin patch (**Figure 4c**). These data suggested that the local adhesion of cadherins at the center of the contact was essential for polarity establishment. Correlatively, signaling from cadherins outside the future luminal area was not sufficient to trigger the formation of apical poles. We concluded that the lumen development originated from the local engagement of cadherins rather than from an integrated signal over the whole contact.

**Figure 4:**
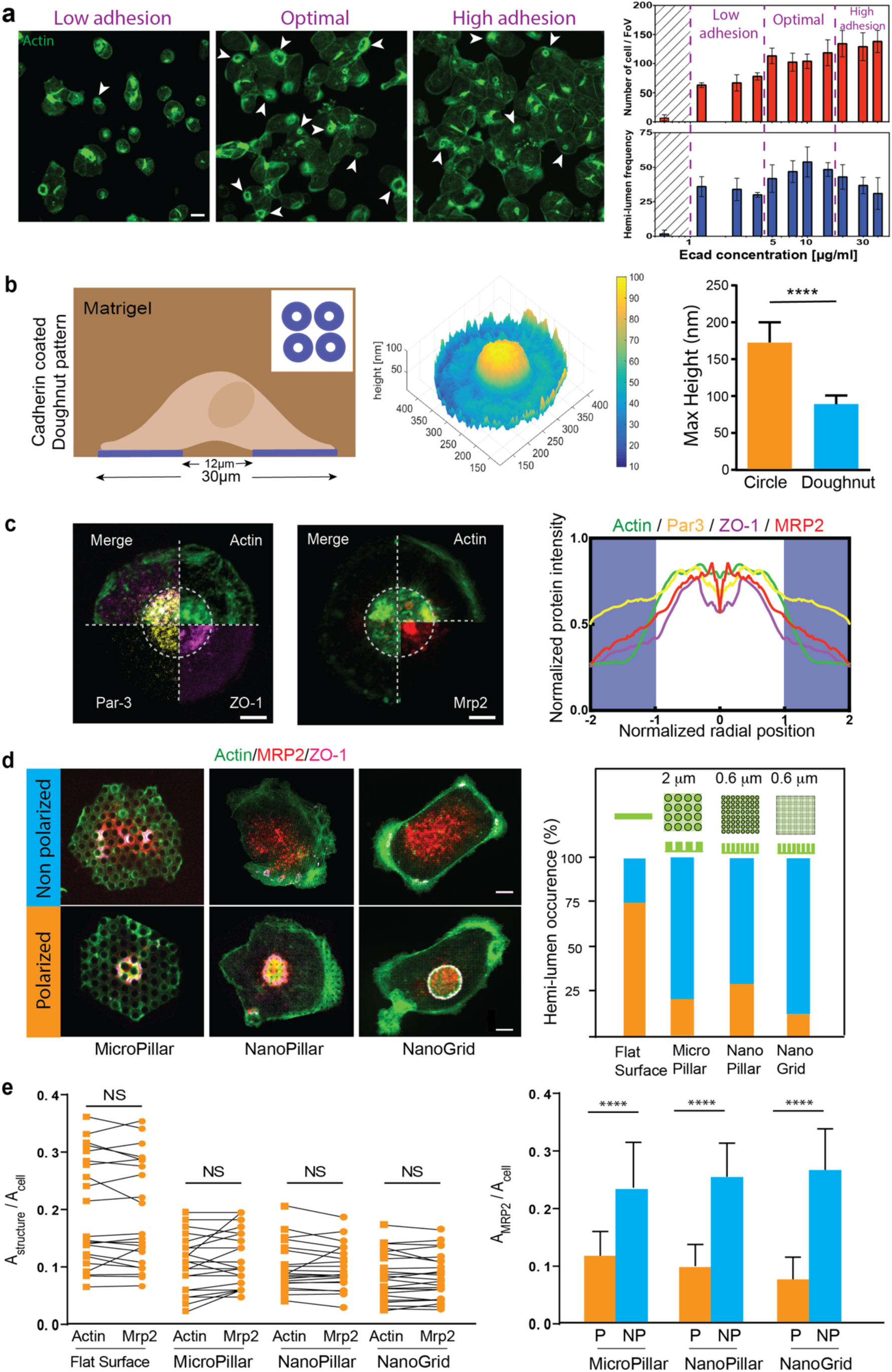
Cadherin-distribution-dependent actin organization is critical for apico-basal polarity establishment. **a,** Low magnification images of primary hepatocytes stained for actin after seeding on different concentrations of E-cadherin. A minimum threshold of E-cadherin concentration is required for the cells to attach, illustrated by the shaded region on the right graph. The number of cells attaching increases with the concentration of E-cadherin above this threshold value. The frequency of hemi-lumen formation reaches an optimum at 10μg/ml E-cadherin, above which the cells preferentially form lumens between each other. White arrows point to representative hemi-lumens. Experiments have been performed on 12 fields of views in 4 independent experiments for each concentration. Scale bar, 10μm. **b, Left panel**: Schematic of the geometry and dimensions of the E-cadherin coated doughnut pattern. **Middle panel**: Representative heat map showing the height of plasma membrane to the coverslip measured from reflective interference contrast microscopy (RICM) image on doughnut pattern of e-cadherin. **Right panel**: Maximum height of plasma membrane quantified from RICM images of primary hepatocytes on E-cadherin circular (n=28) and doughnut (n=14) patterns. ****,p<0.0001. **c,** Representative fluorescence images showing the actin, ZO-1, Par3 (left) and actin, MRP2 (middle) localized on the plasma membrane/E-cadherin interface. Scale bar: 5µm. Quantification of actin, ZO-1, Par3, MRP2 distribution in relation to the position of the non-adhesive region (white) and E-cadherin coated region (Purple), n= 16 for cells stained with actin, ZO-1 and Par3, n=11 for cells stained with actin and MRP2. Despite the creation of a lumen-like structure on doughnut pattern, no specific localization of apical markers has been identified. **d,** Representative immunofluorescence images showing perturbation of Actin, MRP2 and ZO-1 distribution in hepatocytes with either a disorganized central actin phenotype (top, non-polarized NP) or with central actin ring phenotype (below, polarized P) when seeded on micropillar, nanopillar, and nanogrid substrates coated with E-cadherin. Scale bar, 5µm. Quantification of hemi-lumen occurrence in cells on flat surface and on textured substrates as indicated (N= 53, 82, 81 and 100 for flat, micropillar, nanopillar, and nanogrid, respectively). Schematics show the dimension of each texture. The number of cells analyzed was pooled from 4 independent experiments. **e,** Quantification of the ratio of the area of the central actin ring and MRP2 (A_structure_) to the cell area (A_cell_) for polarized primary hepatocytes seeded on flat surface (n=22), micropillar (n=25), nanopillar (n=20) and nanogrid (n=24). Black lines pair the ratio measured for actin and MRP2 in the same hepatocyte. Quantification of the ratio of the area of MRP2 to the cell area of polarized (P) and non-polarized cells (NP) seeding on micropillar, nanopillar and nanogrid (n=24-35), ****, p<0.0001.

We reasoned that the development of actin fibers (P2 phase) and the subsequent diffuse actin patch (P3 phase) from an actin poor contact area (P1 phase) was an essential process in the subsequent self-organization of the apical pole. To test this hypothesis we impaired the development of the fibers while still maintaining the full capacity of the hepatocytes to self-organize. We plated the hepatocytes on a substrate homogeneously coated with E-cad that was studded with small topographical features (comprising of pillars or a grid with dimensions ranging from 2um to 500 nm in width, and 800nm in height, **Material and Methods**). Hepatocytes were able to attach to such substrates, spreading on and in-between the topographical features. Despite the homogeneous E-cadherin coating, all the feature types tested resulted in inhibition of the development of P2 like actin fibers (**Figure 4d**). Actin accumulated around each topographical feature fully coated with E-cadherin.

Upon addition of matrigel, most of the actin failed to restructure and remained largely “clamped” by the topography. This resulted in a drastic reduction of hepatocytes with a polarized phenotype (75% polarization in the absence of features, compared to 20% polarization on micropillars, 25 % on nanopillars, and 15% on nanogrids, **Figure 4d**). The few polarized cells (discriminated by an actin structure surrounded by ZO1 and Par3) exhibited a much smaller apical area (2~20% vs 8~30% for control)(**Figure 4e**). Independent of the apical pole size, MRP2 perfectly overlaid the actin structure. However, in the non-polarized phenotypes, MRP2 was diffusively recruited at the contact with the substrate (**Figure 4d,e).** Our data demonstrated that the self-organizing process of the cell autonomous hepatic polarity development was orchestrated by the ability of the actin cortex to reorganize from a suspended cortical actin structure into a loose actin meshwork at cell-cell contacts. Engagement of integrin signaling at the basal poles leads to the development of actin fibers at the apical pole. The development of the incorrect actin structure (either by physical impairment or by local absence of cadherin adhesion) led to the inhibition of the apical pole development and all subsequent polarity development.

In conclusion, we have established a novel single cell model to investigate the role of of cell-cell junction in apical basal polarity. Our data demonstrate that the development of apical basal polarity in hepatocytes is a largely cell autonomous process, independent of the nature of their epithelial neighbors. Our results strongly suggest that lumen formation can occur between mature and immature hepatocytes during development. Hepatic polarity appears as an emergent property induced by the spatially segregated contact of cadherin and ECM adhesion without any need for collective cellular response. The structural and mechanical properties of the actin cortex at the lateral contact acts as a switch triggering the development of the apical pole. The density and spatial distribution of cadherins at the initial cell-cell junction largely regulates the development of an apical actin cortex that in turn drives the polarized organization of the whole cell including that of the Golgi apparatus.

Our reductionist approach demonstrate that single hepatocytes can be fooled into a polarized state by artificial microniches and thus constitutes, as far as bile secretion is concerned, the first realization of a single cell liver.

## Methods

### Microwell fabrication

Microwells with dimensions of 25µm in diameter and 25µm in height were fabricated using an established method (*29*). The functionalization of the microwell top, side, and bottom surfaces was achieved by coating with 10µg/ml fibronectin (Sigma, P1141) for 1hour, followed by flipping into a fibronectin coated coverslip to passivate the new-top surface with a solution of 0.2% pluronic acid (Sigma, P2443-250G).

### Generation of Micropatterned substrates

The 2D patterns were generated by microserigraphy method (*30*). 100µl of 10 µg/ml fibronectin or E-Cadherin (R&D System, 8875-EC) was applied to a 2X2 cm NoA 74 membrane on a polymer bottom dish (ibidi, 81156), and incubated overnight at 4°C. The membrane was peeled off right before usage and the dish was treated with 0.2% pluronic acid for 30min at RT.

Employing the Alveole PRIMO system, E-Cadherin coated doughnut patterns were produced as recommended by the vendor. 10µl of 100µg/ml E-Cadherin solution was applied to each PDMS stencil, incubated for 2 hours at room temperature before rinsing with PBS 3 times. The dish was then treated with 0.2% pluronic acid for 30min.

### Topographical obstacles fabrication

Replicas of silicon molds containing the different features (750nm in height) was made by double-casting PDMS (mixed at 10:1 base and curing agent, Sylgard184, Dow Corning) cured at 80°C for 3 hours, passivated overnight at low pressure with a solution of Trichloro(1H,1H,2H,2H-perfluorooctyl)-silane (Sigma, 448931).

The textured substrates were generated by UV curing (6 min, 185 and 253nm, 30mW/cm2, UVO Cleaner 342-220, Jelight) a drop of low refractive index polymer premix (MY134, MyPolymers), sandwiched between a glass coverslip and the PDMS mold, and immersed in water. After peeling off the mold, the features were coated overnight with 10μg/ml E-cadherin solution (RnD, 8875-EC-50) in PBS at 4°C and washed twice with PBS before cell seeding.

### hiPSC differentiation, seeding and culturing

hiPSC-derived hepatic progenitor or hepatocyte-like cells were generated using an established protocol (*18, 19, 31*). To generate heterodoublets of iPSC-derived hepatocytes at mature and immature stages, hepatocyte-like cells after 25 days of differentiation were detached and suspended in Hepatozyme medium (Thermo, 17705021), supplemented with Oncostatin M 0.01 mg/ml (Bio-Techne, 295-OM-050) and Hepatocyte Growth Factor 0.05 mg/ml (Peprotech, 100-39-100) to reach a final cell density of 0.5 * 10^6^ cells/ml. Approximately 1 ml cell suspension was then pipetted onto microwells in a 35mm glass bottom dish and placed in an incubator for at least 2 hours to allow cell attachment. Extra cells that were not trapped in the wells were removed by rinsing the dish with PBS buffer. The system was then replenished with fresh culture medium. Cells were left in 5% CO_2_ at 37°C and 95% humidity for 1 day to develop polarity.

### Hepatocyte isolation, seeding and culturing

Hepatocytes were isolated from male Wistar rats by a two-step in situ collagenase perfusion method, as previously published (*32*). Animals were handled according to the IACUC protocol approved by the IACUC committee of the National University of Singapore. With a yield of >10^8^ cells/rat, hepatocyte viability was tested to be >90% by Trypan Blue exclusion assay.

In order to co-culture primary rat hepatocytes with another epithelial cell line, e.g. MDCK, EpH4, Caco2 in a microwell array, freshly isolated rat hepatocytes (0.5 million) were seeded onto the microwell in the glass bottom dish and cultured in 2 ml of William’s E culture medium supplemented with 2 mM L-Glutamine, 1 mg/ml BSA, 0.3 μg/ml of insulin, 100 nM dexamethasone, 50 μg/ml linoleic acid, 100 units/ml penicillin, and 100 mg/ml streptomycin (Sigma-Aldrich). After 1 hour incubation, the floating hepatocytes were removed by washing with PBS buffer and culture medium were replenished. 0.5 million MDCK cells expressing histone-GFP (generous gift from Dr Benoit Ladoux, Institut Jacques Monod, Paris), EpH4 or Caco2 cells stained with CellTracker^TM^ Green CMFDA Dye (ThermoFisher, C2925) following manufacturer’s instruction were subsequently detached and seeded into the microniches. After 1 hour incubation, excess cells were removed and culture medium were replenished. The system was left in incubator for 24 hours to develop polarity.

For micropatterning experiments, 0.5 million rat hepatocytes were added onto E-Cadherin coated micropatterns in a 35mm glass bottom dish and cultured in 2ml of William’s E medium with all seven supplements as described before. Cells were incubated with 5% CO_2_ at 37°C and 95% humidity. After a 3-hour incubation, the system was rinsed with PBS medium to remove hepatocytes that did not attach to the micropatterns. The petri dish was subsequently replenished with fresh culture medium. 3-hours later, the culture medium was replaced by medium supplemented with 6% matrigel. The matrigel was handled according to the protocol as described previously[Martın-Belmonte, 2013]. The system was then left in incubator for 24 hours.

### Pharmacological treatment

To inhibit actomyosin contractility or block bile acid synthesis, culture medium supplemented with blebbistatin (50μM in DMSO; Merck, 203390) or ketoconazole (10µM in DMSO; Sigma, K1003) was administered 7 hours after matrigel overlay until cell fixation. To stimulate bile acid secretion, Ursodeoxycholic acid (UDCA, 50µM in DMSO; Sigma, U5127) was added at the same time as the matrigel overlay.

### Immunostaining and image acquisition

Cells were fixed with 4% paraformaldehyde (PFA) for 30 minutes at 37°C. After fixation, the cells were rinsed with PBS and permeabilized for 30 min in PBST (0.1% Triton-X diluted in TBS). Permeabilized cells were blocked with 5% BSA diluted in PBS for 4 h at 4°C and incubated overnight with pan-Cadherin antibody (Sigma, C1821, 1:500), MRP2 antibody (Sigma, M8316, 1:200), Par-3 antibody (Millipore, 07-330, 1:200), ZO-1 antibody (Life Technology, 61-7300,1:100), ZO-2 antibody (ThermoFisher, 38-9100, 1:100), Claudin-1 antibody (Invitro, 717800, 1:100), Claudin-3 antibody(Abcam, ab15102, 1:40), Occludin antibody (Invitrogen, 711500 1:200), MyosinIIA antibody (Sigma, M8064, 1:200), Grasp65 antibody(Abcam, ab102645, 1:200) at 4°C as instructed in the manufacturer’s protocol. After rinsing with PBS, cells were incubated with secondary antibodies (Alexa Fluor 546 Donkey Anti-Rabbit IgG, A10040 and Alexa Fluor 647 Donkey Anti-Mouse IgG, A-31571, Life Technologies, 1:200) and Alexa Fluor 488 Phalloidin (Life Technologies, A12379, 1:200) or ATTO-565 Phalloidin (Sigma 94072, 1:500) for 1 h in dark at room temperature. After rinsing with PBS again and incubation with DAPI (Sigma, D9564), cells were mounted in mounting medium (DAKO, S3023). 3D stacks of confocal images were acquired with 60X NA1.3 water lens on a Nikon Eclipse Ti Microscope equipped with Yokogawa CSU-X1 spinning disc unit. Structured Illumination Microscopy images was acquired on the same microscope equipped with Live-SR module (https://www.cairn-research.co.uk/product/live-sr/). The cells were chosen purely based on criteria of cell adhesion. Typically, more than 70% of patterns contained single cells that occupied the entire pattern, and these were selected for imaging.

### RICM analysis

RICM analysis was performed considering the theory of partial coherent light, following the description of cell adhesion analyses reported in Limozin and Sengupta (*25*). Relative heights were reconstructed using the intensity-height relation

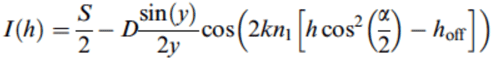

where

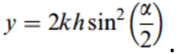

k = 2π/λ is the wave vector for the illumination light for a wavelength λ = 546 ± 10 nm, n1 = 1.335 is the refractive index of the outer buffer, S and D are the sum and difference of the maximal and minimal intensity in the experimental fringe pattern, respectively, and h_off_ is a phase shift arising from the reflection at different interfaces.

The illumination numerical aperture (INA), which is given by the half-angle of the cone of illumination, α, was set to a maximum value to minimize the depth of focus and thereby to avoid reflections from organelles or other intracellular structures. The measured INA amounted to INA = n_1_ sin(α) = 0.73. Cell contact areas of constant dark intensity were considered ‘adhered’ and of clostest proximity to the substrate. These areas served as starting point for the reconstruction of relative membrane heights.

Data were analyzed using self-written routines in Matlab (version 9.3 (R2017b), The MathWorks, Inc. MA, USA) and FIJI (version 1.52s, Rasband, W.S., NIH, Bethesda, MD, USA).

### Image analysis

To analyze the relative position of each protein, a homemade program was written with Matlab (MathWorks, Natick, Mass). The position of the lumen center was determined by using fit-circle function implemented in Matlab with a radius range lower than the cell size on the thresholded actin image by Otsu’s method acquired at the membrane/substrate interface. The radial profile was then performed on every channel. The intensity profile for each cell was then aligned by normalizing the distance between the centre of the lumen and the outer point of the actin ring. This outer point was determined by finding the maximum of the second derivative of the actin intensity profile. As there was no ring formed on E-cadherin doughnut pattern, the distance was normalized between the centre of the pattern and the inner border of the adhesive area. The relative distance of the different ring was then calculated by measuring the distance between the peaks of the average curve of each staining.

To assess Golgi localization in relation to lumen position, the degree of overlay of these two structures was measured. Z direction maximum intensity projection was applied to all the stacks containing Grasp65 signal to extract Golgi structure, while projection of selected frames of Phalloidin staining at the cell/substrate interface was used to extract lumen localization. ROIs of Golgi and lumen structures were created by thresholding the corresponding Z-projection images. The ratio of the number of pixels of the intersection over that of actin mask was finally used to describe the degree of overlaying.

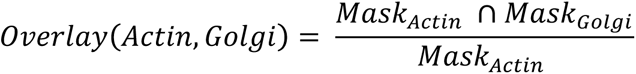

To measure the size and circularity of hemi-lumens and cells, 3 frames of phalloidin staining images at the hepatocyte/substrate interface were selected and reconstructed using maximum intensity projection. The contour of the hemi-lumen and cells was drawn manually based on F-actin signal using ImageJ. The Area and Circularity were measured with the ImageJ measurement plugin. A circularity of 1.0 indicates a perfect circle. As the value approaches 0, it indicates an elongated polygon.

To evaluate the MRP2 distribution and actin structure size for hepatocytes cultured on textured substrate, selected frames imaged at the hepatocyte/substrate interface were reconstructed using maximum intensity projection. The areas of MRP2, actin structure and cells were then manually measured using ImageJ.

Statistical analysis was performed using GraphPad Prism 6 (https://www.graphpad.com/scientific-software/prism/). The statistical significance between two groups was analyzed by Unpaired Student’s t-test unless otherwise stated. In all cases, a P value of less than 0.05 was considered statistically significant and P value is specified in each captions.

## Supplementary Figures

**Supplementary Figure 1:**
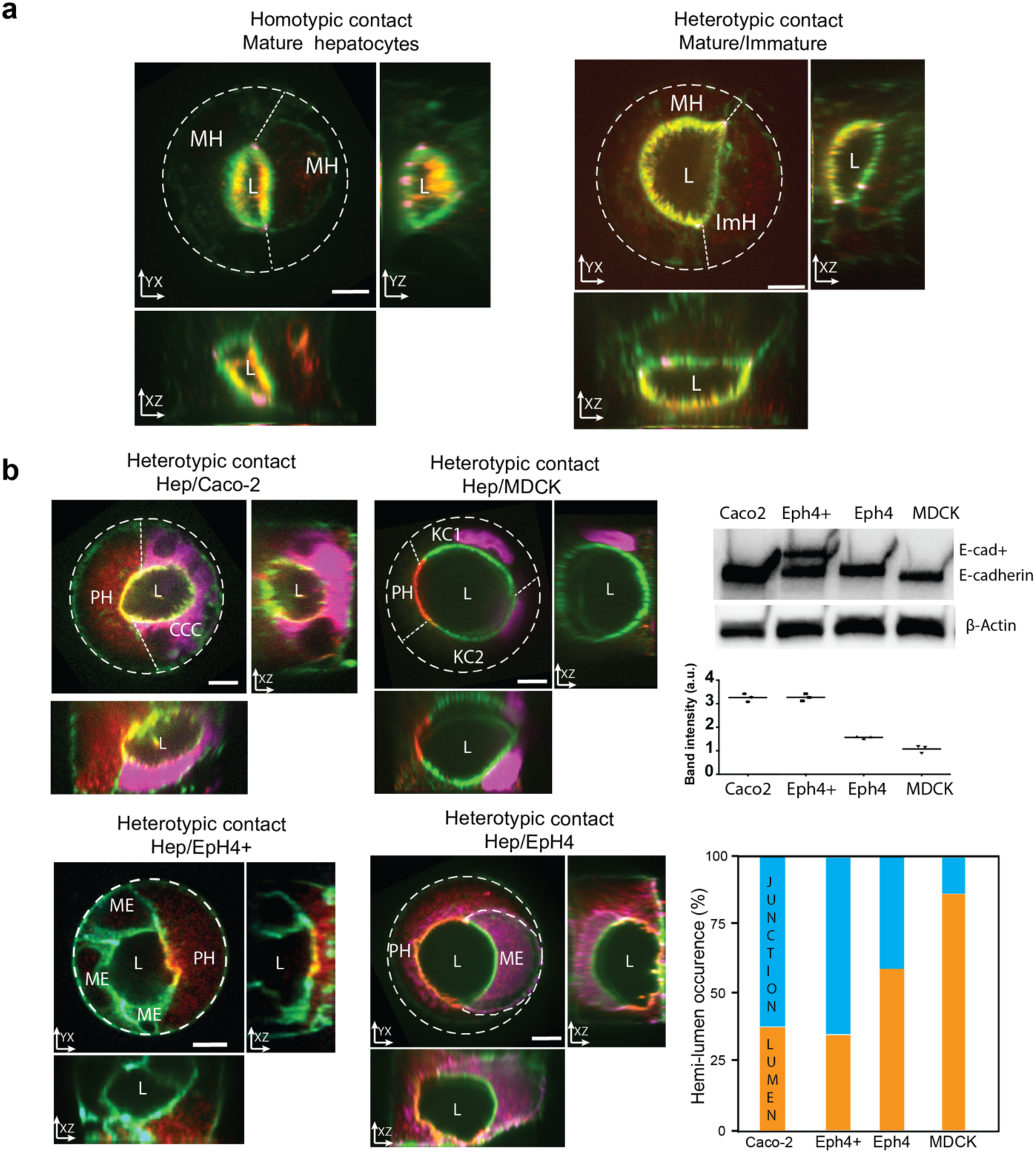
Orthogonal views of heterodoublets forming lumen-like structures. **a,** Orthogonal views of representative images for Figure 1 showing lumens formed between mature (MH) and immature (ImH) hiPSC derived hepatocyte. MRP2 (red) is exclusively recruited to the apical domain of mature hepatocytes. Dash lines indicate the cell-cell contacts. **b**, Top and side views of heterodoublets between primary rat hepatocytes (PH) and different epithelial cell lines (MDCK, Eph4, EPH4+Ecad, Caco2). Discrimination between cell types was performed using prestaining before the formation of the doublets. Caco2 and Eph4 cell lines were pre stained using cell tracker, Eph4+ expressed E-cad GFP and Actin is in green, MDCK expressed H2B-GFP. E-cadherin levels in each cell lines were assessed by three indepednetn westernblot. The hemi-lumen occurrence is inversely correlated to expression levels of E-cadherin. Scale bar, 5μm. N_Caco2_ = 29 (38%), N_Eph4+_ = 20 (35%), N_Eph4_ = 56 (59%), N_MDCK_ = 53 (87%),

**Supplementary Figure 2:**
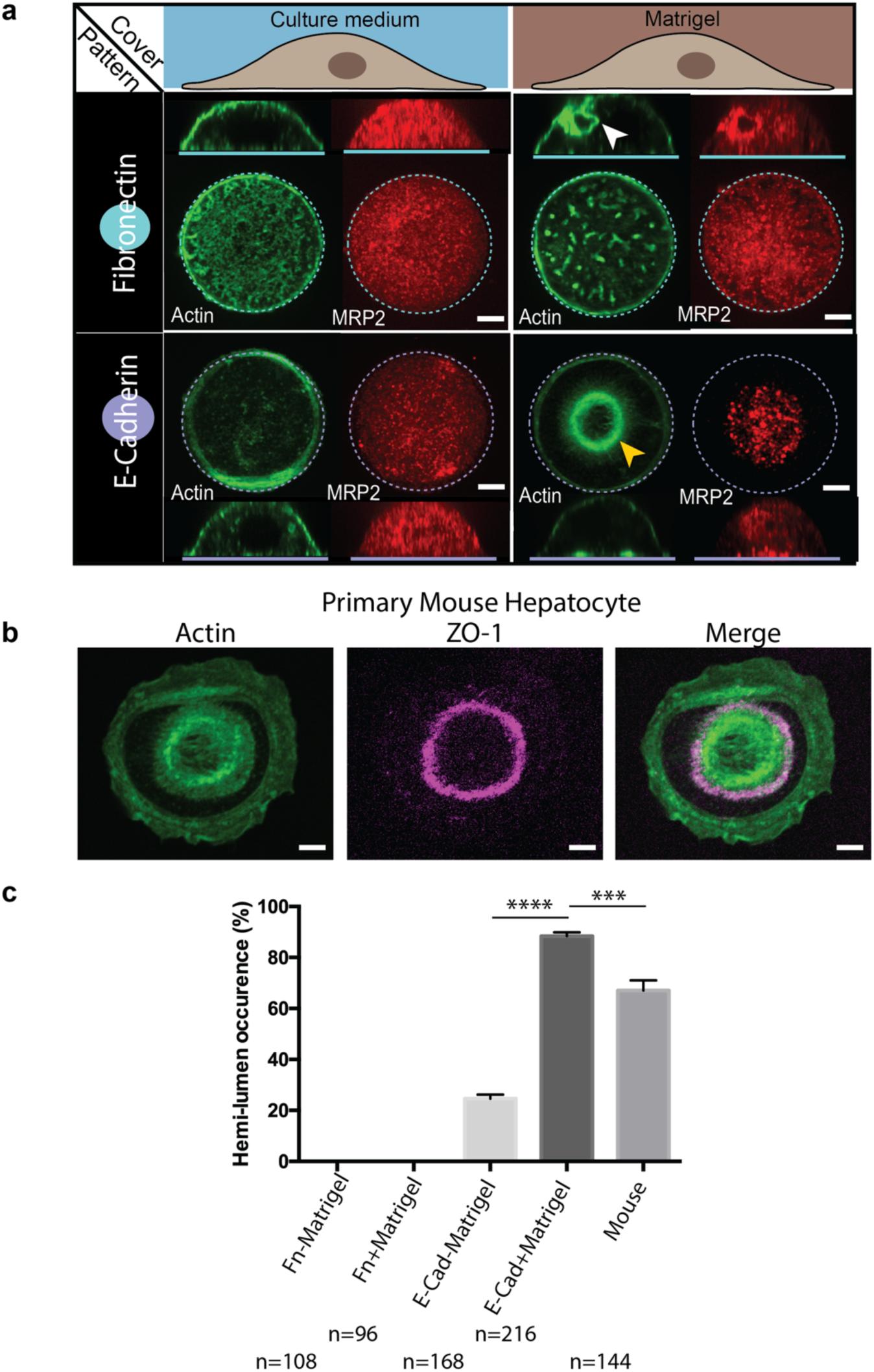
Combination of Cadherin and ECM signalling are necessary to form hemi-lumen in large proportion of hepatocytes. **a,** Representative immunostaining images showing the top and side view of single hepatocytes cultured in four distinct microenvironments as indicated by the diagram. Scale bar, 5μm. **b,** Representative image showing that primary mouse hepatocyte are capable of forming similar actin (Green) structure with ZO-1(purple) exclusively recruited to the contour as rat hepatocyte when cultured on E-Cadherin coated island and and overlayed with 6% matrigel. Scale bar, 5μm. **c,** Fraction of cells displaying central actin structure and polarity phenotype in conditions described in a, b. The data of rat and mouse hepatocyte is collected from 3 and 2 independent experiments respectively. ***, p value < 0.001, ****, p value < 0.0001.

**Supplementary Figure 3:**
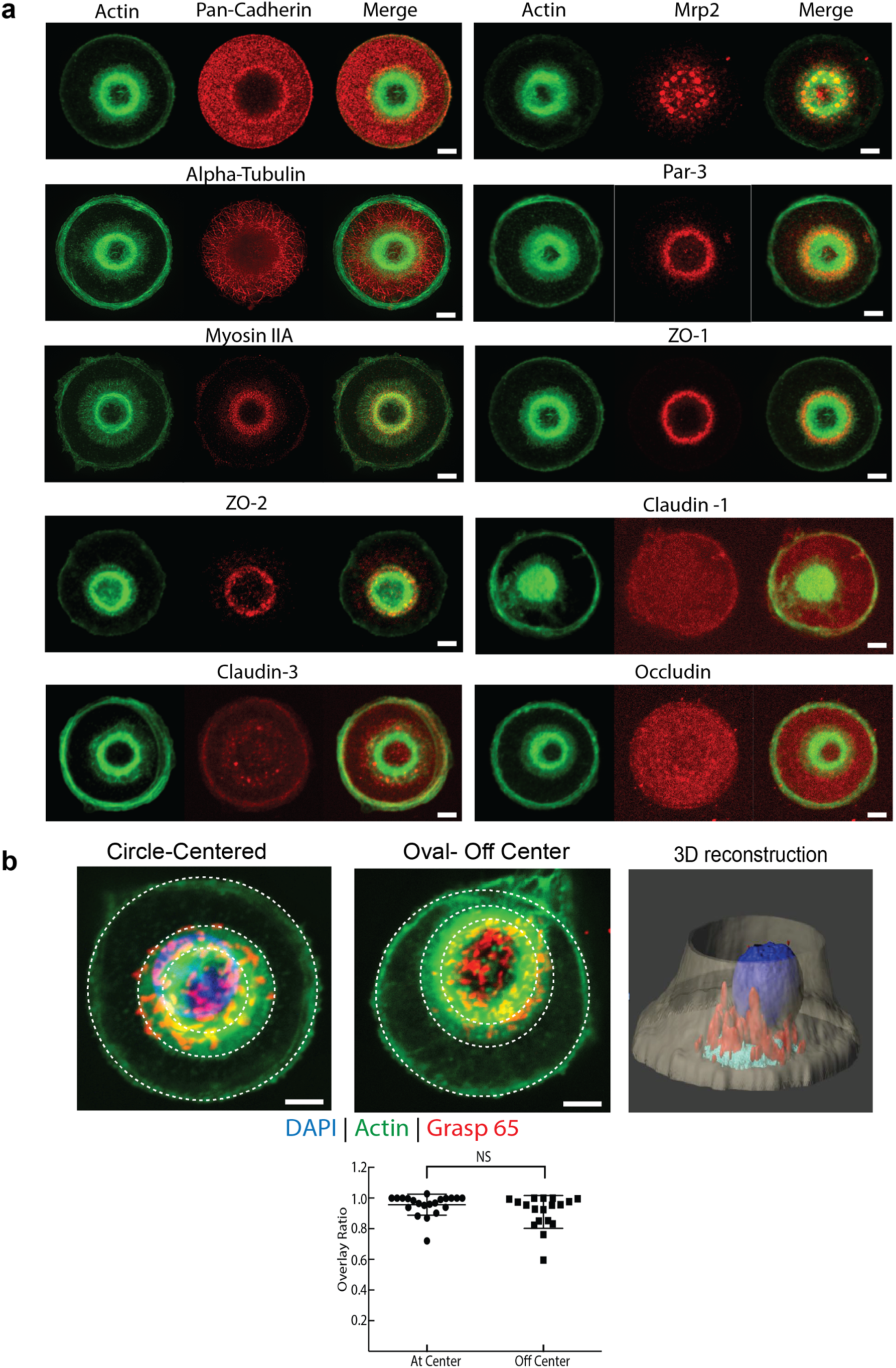
The distribution of polarity markers protein and organelles in hemi-lumen. **a,** Split color images for Fig. 2b. Representative immunofluorescence images of ZO-2, Claudin-1, Occludin and Alpha-Tubulin that are not presented on Fig 2b. Scale bar, 5μm. **b,** Representative images showing relative location of actin structure (green) and maximum Z projection of Golgi staining (by Grasp65 in red) in the cases when the hemi-lumens at the centre or side of cell/substrate interface. 3D reconstruction (middle) of image stack for left panel showing the Golgi(red) is situated right above lumen area (Cyan). Plots (right) showing the projection of Golgi structure highly overlapped with lumen area. We do not observe any significant difference between both cases of lumen location. n=21 for centered lumen at centre, n=19 for off-centered lumen.

**Supplementary Figure 4:**
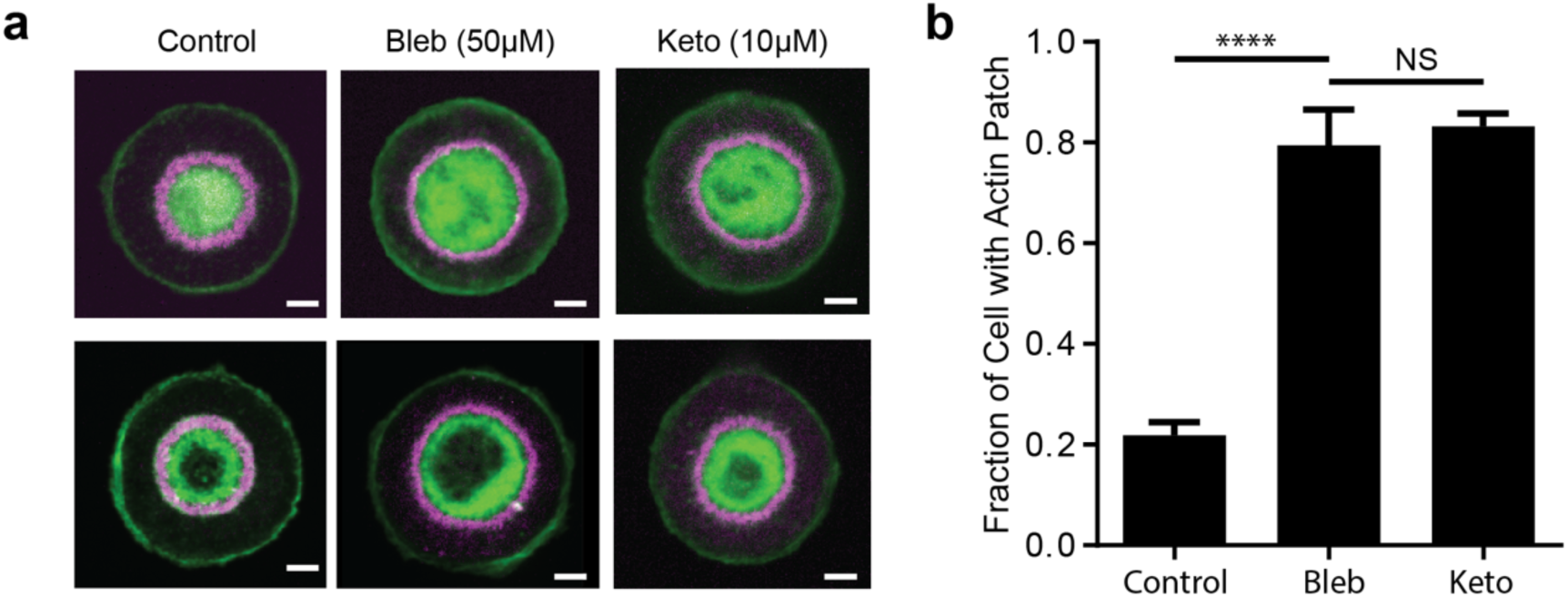
Inhibition of actomyosin contractility or of bile acid synthesis hinders the reorganization of the actin structures from dense patches to rings. **a,** Representative images of actin (green) and ZO-1 (purple) for both patch (top) and ring (bottom) morphologies in control, 50μM blebbistatin (Blebb) and 10μM ketoconazole (Keto) treated hepatocytes. Scale bar: 5μm **b,** Fraction of cells with dense patch actin structures at steady state is significantly increased after blebbistatin or ketoconazole treatments. Bar chart shows the mean±SD from 3 independent experiments (n=146, n=129, and n=108 cells in control, Bleb (Blebbistatin) and Keto (Ketoconasol) group respectively). ****, p<0.0001, NS, not significant. Two-sided unpaired t-test was performed to calculate p value.

**Supplementary Figure 5:**
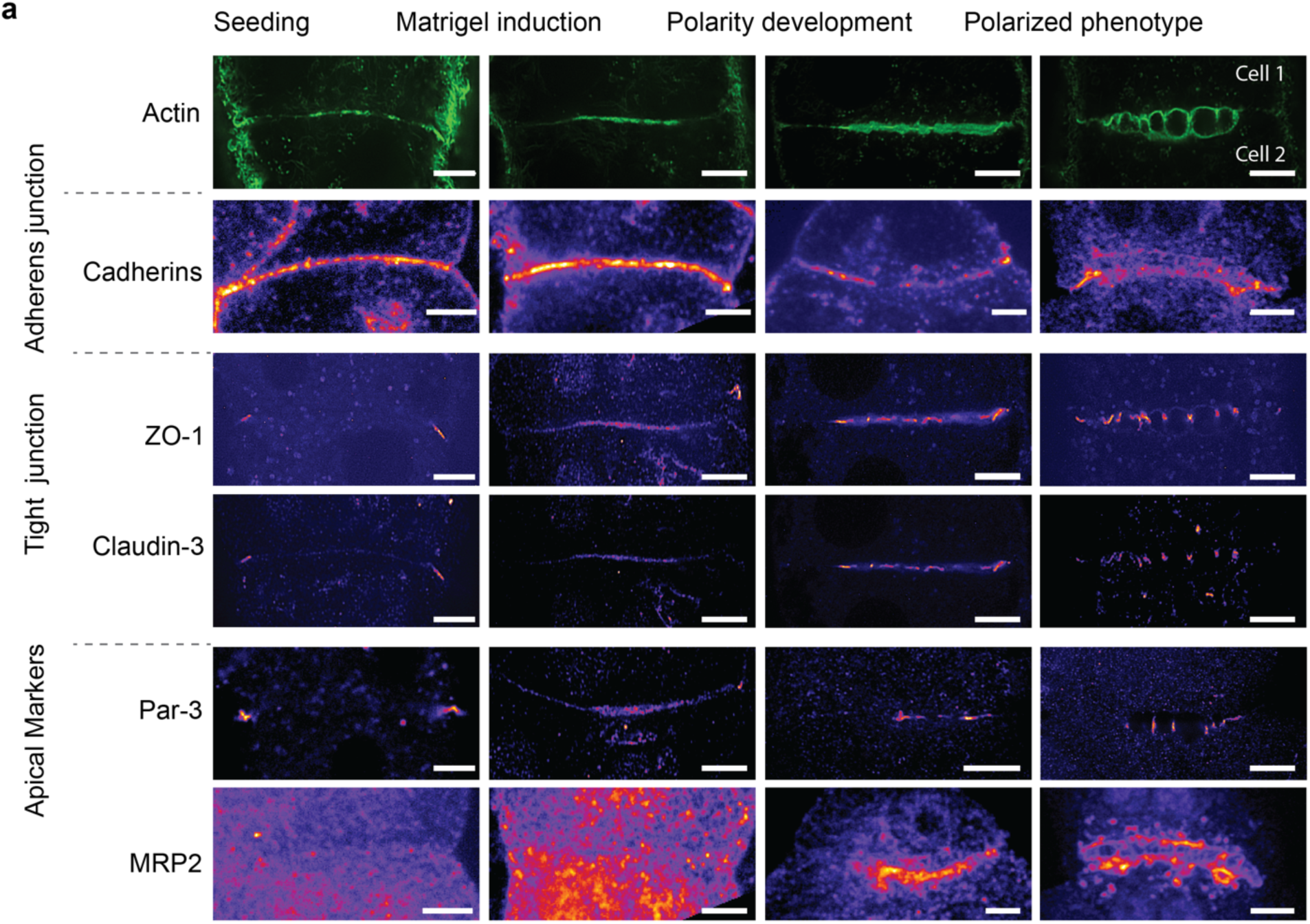
Evolution of the distribution of polarity markers around the cell-cell interface during *in vivo* polarisation. Representative structure illumination microscopy (SIM) images of actin and hepatic polarity related proteins for bile canaliculi formed a lumen between two primary hepatocytes at different stages of polarity development. Actin, cadherin and ZO1 and Claudin 3 images are obtained from quadruple immunostaining of the cells. Par 3 and MRP2 images are obtained from cells with similar actin morphologies.

**Supplementary Figure 6:**
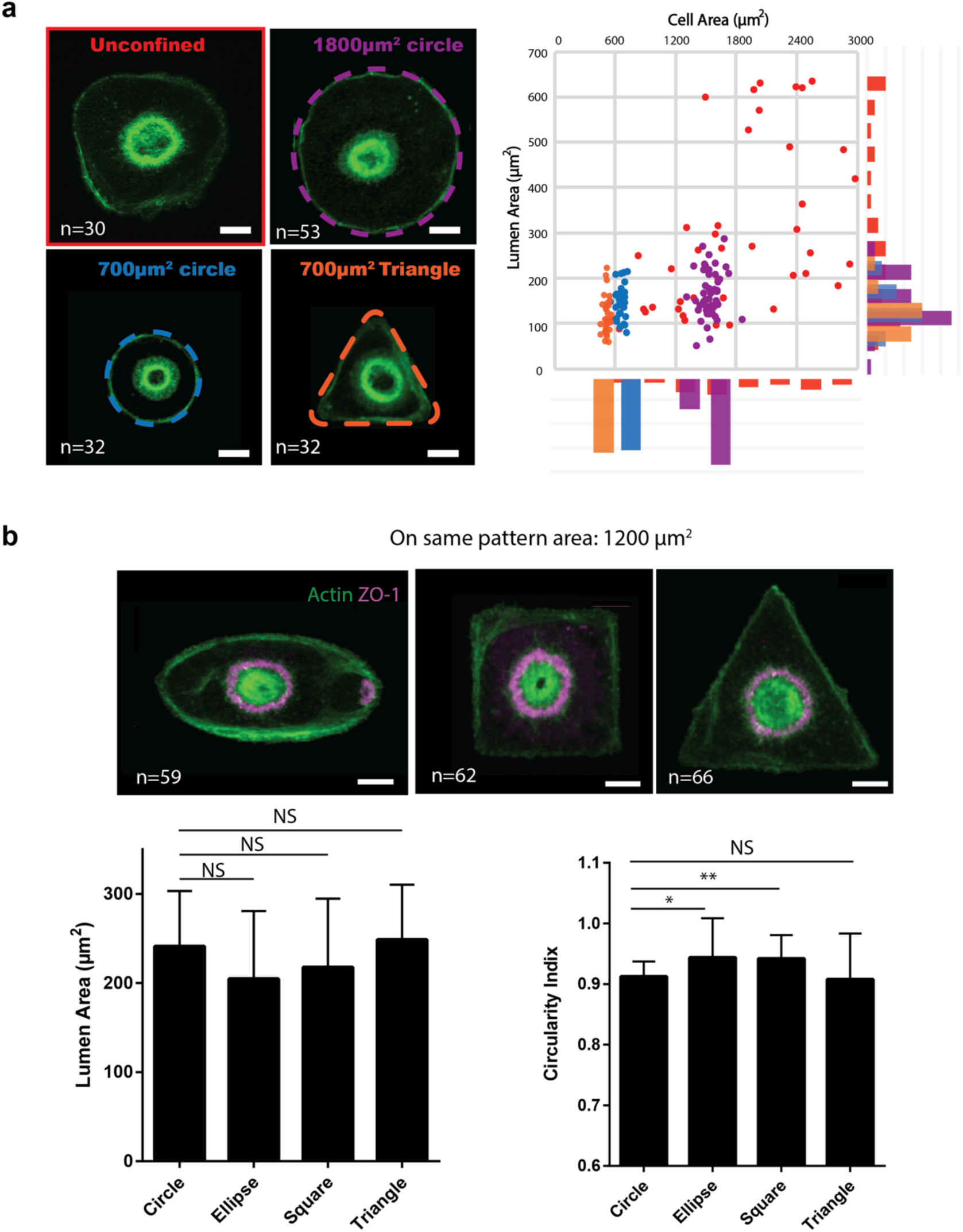
The hemi-lumen size and shape are relatively independent of the cell spreading area and geometry. **a, Left panel:** representative images of actin (green) showing the sizes and shapes of hemi-lumens formed by hepatocytes spreading on E-Cadherin coated surface with different areas. The patterns are outlined by coloured dashed lines. Scale bar:5μm. **Right panel**: Distribution of lumens area vs and cell area. Data from cells on different spreading areas or geometries are color-coded corresponding to the color of the pattern outlines depicted on the left panel. The data are pooled from 3 independent experiments. **b, Top:** typical central actin and ZO-1 staining of hepatocyte seeding on elliptic, square and triangular pattern with area of 1200 µm^2^. **bottom**: bar charts showing the lumen area (left) and circularity index (right) of hemi-lumen in cells confined on different cell geometry. NS, not significant, **, p<0,01, *, p<0,05.

**Supplementary Figure 7:**
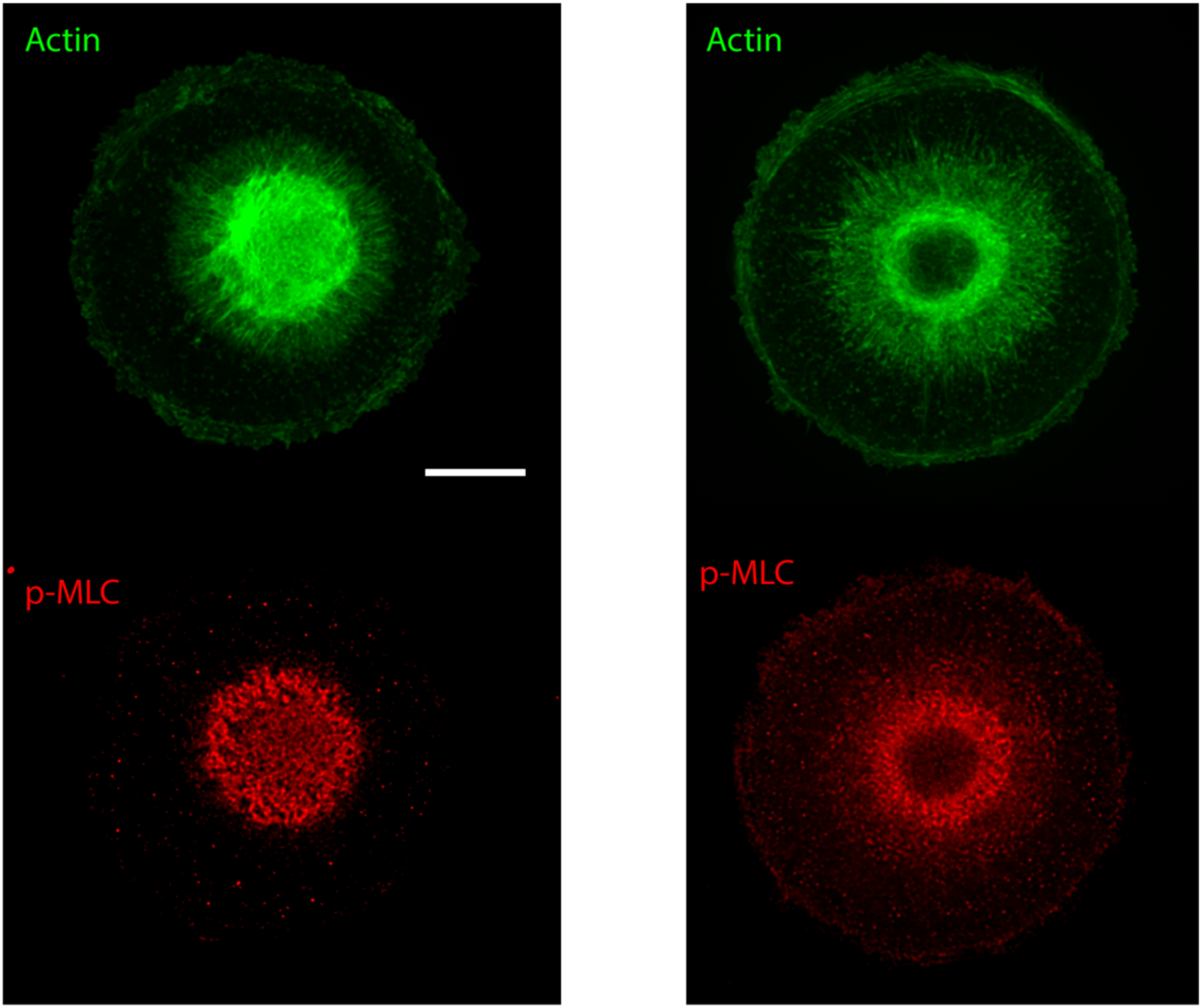
Actin and Myosin organisation during hemi-lumen formation imaged with structured illumination microscopy (SIM). **Left**,: Superresolved image of the Actin patch of phase P4 (Green) and phospho Myosin light chain localization. The radial fibers around the patch are largely deprived of p-MLC while the dense actin structure in the centre display myosin minifilaments. **Right:** Actin ring in phase P5. The ring concentrate most of the actomyosin complexes, indicating a local contractility. Scale bar = 10μm

### Caption Supplementary movies

**Supplementary Movie 1:** live RICM imaging of the hemi-lumen in control conditions. Note the central ring that moves radially indicating a dome shape structure of about 200 nm height on top of the coverslip.

**Supplementary Movie 2:** live RICM imaging of the hemi-lumen under Ketokonazole 10 uM. The inhibition of bile secretion leads to a large reduction of vertical fluctuation (amplitude and frequency). The fluctuations are no longer radial (lumen pulsating) but rather diffused and lateral, indicating a mere membrane fluctuation.

**Supplementary Movie 3:** live RICM imaging of the hemi-lumen in UDCA (40uM) treated cells. This treatment stimulates bile salt secretion. Multiple interference rings in the center of the lumen indicates a much-inflated geometry compared to control case.

**Supplementary Movie 4:** live RICM imaging of the hemi-lumen on doughnut patterns in control conditions. The central fluctuations are fully random compared to disc patterns. It indicates the absence of coordinated pulsations that are replaced by simple fluctuations of the free membrane.

